# Power Analysis of Single Cell RNA-Sequencing Experiments

**DOI:** 10.1101/073692

**Authors:** Valentine Svensson, Kedar Nath Natarajan, Lam-Ha Ly, Ricardo J Miragaia, Charlotte Labalette, Iain C Macaulay, Ana Cvejic, Sarah A Teichmann

## Abstract

High-throughput single cell RNA sequencing (scRNA-seq) has become an established and powerful method to investigate transcriptomic cell-to-cell variation, and has revealed new cell types, and new insights into developmental process and stochasticity in gene expression. There are now several published scRNA-seq protocols, which all sequence transcriptomes from a minute amount of starting material. Therefore, a key question is how these methods compare in terms of sensitivity of detection of mRNA molecules, and accuracy of quantification of gene expression. Here, we assessed the sensitivity and accuracy of many published data sets based on standardized spike-ins with a uniform raw data processing pipeline. We developed a flexible and fast UMI counting tool (https://github.com/vals/umis) which is compatible with all UMI based protocols. This allowed us to relate these parameters to sequencing depth, and discuss the trade offs between the different methods. To confirm our results, we performed experiments on cells from the same population using three different protocols. We also investigated the effect of RNA degradation on spike-in molecules, and the average efficiency of scRNA-seq on spike-in molecules *versus* endogenous RNAs.

## Introduction

Recently, there has been an explosion in the development of protocols for RNA-sequencing of individual cells (single cell RNA-sequencing, scRNA-seq)^1,2^ which make use of different cell capture and DNA amplification strategies, techniques to combat biases, and liquid handling platforms. Due to the tiny amount of starting material, a considerable amount of amplification is an integral step of all of these protocols. Consequently, it is important to assess the sensitivity and accuracy of the protocols in terms of numbers of RNA molecules detected. An objective strategy to assess the technical variability in these methods is to add exogenous spike-in RNA of known abundances to the individual cell samples. In this study, we assessed the performance of several published scRNA-seq methods according to their ability to quantify the expression of spike-ins of known concentrations. An ideal method is both sensitive and accurate, as well as cheap, where cost is reflected in sequencing depth.

We define sensitivity as the minimum number of input RNA molecules required for a spike-in to be detected as expressed, and accuracy as the closeness of estimated relative abundances to ground truth (known relative abundances of molecules). High sensitivity permits detection of very lowly expressed genes. High accuracy implies that detected differences in expression reflect true biological differences in mRNA abundance across cells, rather than technical factors.

The spike-in collections designed by the ERCC (External RNA Controls Consortium)^3^ consist of a set of 92 RNA sequences of varying length and GC content. They are used in a mix at known concentrations representing 22 abundance levels that are spaced one fold change apart from each other (Supplementary Figure 1). Previously, such spike-ins have been applied to assess standard RNA-sequencing protocol reproducibility^4^, and to assess performance of differential expression tests in RNA-sequencing data^5^. In the context of single cell RNA-sequencing protocols, ERCC spike-ins were first published as part of the description of the CEL-seq protocol^6^.

Here, we exploit the fact that spike-ins provide us with a means to calculate a technical sensitivity and accuracy for each protocol using different platforms in a way that is comparable and independent of the biological system under investigation (Figure 1A-B). Using knowledge of the input molecules of spike-ins, we can calculate the lower detection limits in number of molecules for each experiment independent of biological differences in systems which were investigated (Figure 1C), and compare this to the overall sequencing depth, giving us the sensitivity at a given depth in number of reads for the various protocols. The ERCC spike-ins also provide a direct way to assess accuracy: we compared the stated input concentration of molecules to the measured expression levels across detected spike-ins (Figure 1D). Thus we obtain a unified framework for comparing sensitivity and accuracy of the various protocols at different sequencing depths.

**Figure 1.**
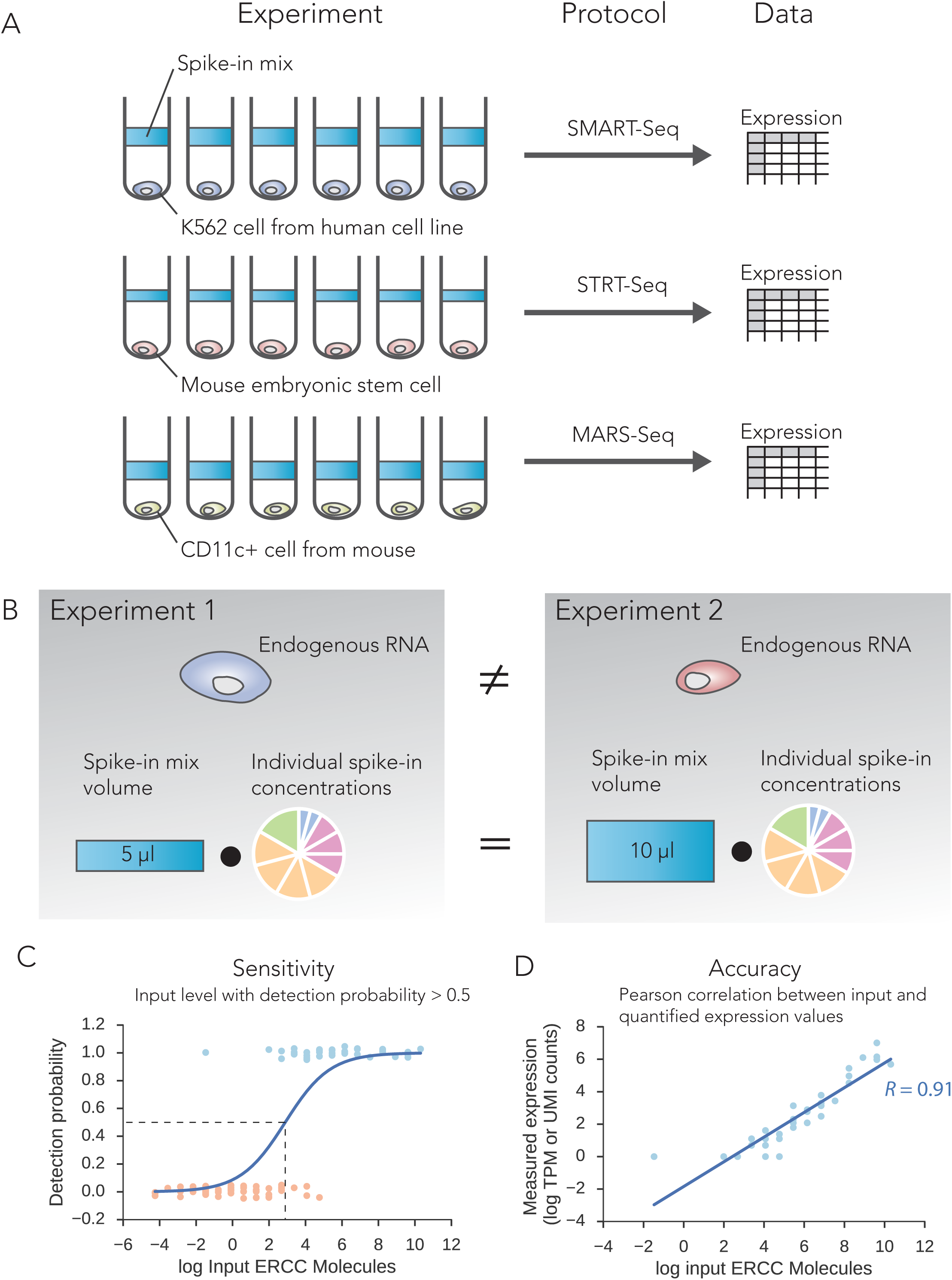
Overview of protocol comparison strategy. **(A)** The data we use are from different protocols and investigate diverse cell types. **(B)** Comparing protocols by looking at properties relating to the cells would be distorted by the diverse cell types involved. Since the same standard spike-in mix has been used in all of them, albeit at different concentrations, we can base our assessment on these synthetic RNA molecules. We define two global technical performance metrics based on these: **(C)** Spike-in sensitivity: the number of spike-in molecules which need to be present in a sample before there is at least 50% chance of detecting them. This is inferred by logistic regression. **(D)** Spike-in quantification accuracy: How well preserved the log-linear relation between input spike-ins is when quantifying the measured expression. We formulate this as the Pearson correlation between input molecules and output expression.

## Results

We retrieved published scRNA-seq data sets which had used ERCC spike-ins for quality control or normalization. We analysed 15 distinct experimental protocols encompassing 28 single-cell studies, of which 17 used a traditional whole-transcript coverage based strategy for measuring expression levels and 11 used strategies based on unique molecular identifiers (UMI’s) for digital quantification of transcripts (Table 1). In addition, we generate our own “dedicated” batch-matched mESC experiment across two replicates. In first batch, we perform SMARTer and Smart-Seq2 on the Fluidigm C1 platform using mES cells and for second batch, we perform SMARTer, Smart-Seq2 and STRT-Seq on the Fluidigm C1. For effective comparison with droplet based technologies, we generate a high throughput dataset on 10X Chromium technology using ERCC spike-ins and control RNA. All coverage based data sets had been sequenced using Illumina paired end sequencing, with read lengths between 75 and 150 base pairs. In total 18,123 publicly available samples were analyzed from about 30×10^9^ sequencing reads.

For each data set, the concentration of mixed spike-ins was noted either through the original study or communication with the original authors, as well as the volume into which they were diluted. From this, we can calculate the number of spike-in RNA molecules from each abundance level that were added to the individual cell samples. This allowed us to compare the samples from the different experimental data sets on the same scale.

### Technical quantification accuracy of different protocols

Above, we compare published data sets in terms of their sensitivity in detecting lowly abundant molecules. How do the various methods compare in their accuracy of quantification of expression levels? We used the 92 ERCC spike-in RNAs to assess the accuracies of the protocols. Each of the 22 abundance levels in the spike-in input-material is two-fold higher than the previous abundance level. We could therefore quantify the accuracy of an experiment by looking at the Pearson correlation between estimated expression levels, and actual concentration of input RNA molecules (ground truth). For each individual cell or sample, we computed the Pearson product-moment correlation coefficient (*R*) between log transformed values for estimated expression and input concentration (Figure 1D). We then compared the values across protocols (Figure 2A). Compared to bulk RNA sequencing, accuracy is consistently lower for scRNA-seq protocols. However, overall, the accuracies are still remarkably high, and rarely do individual samples have a Pearson correlation lower than 0.6. Some protocols exhibit tails towards the lower end of accuracy, which might be indicative of variable success rates of protocols.

**Figure 2.**
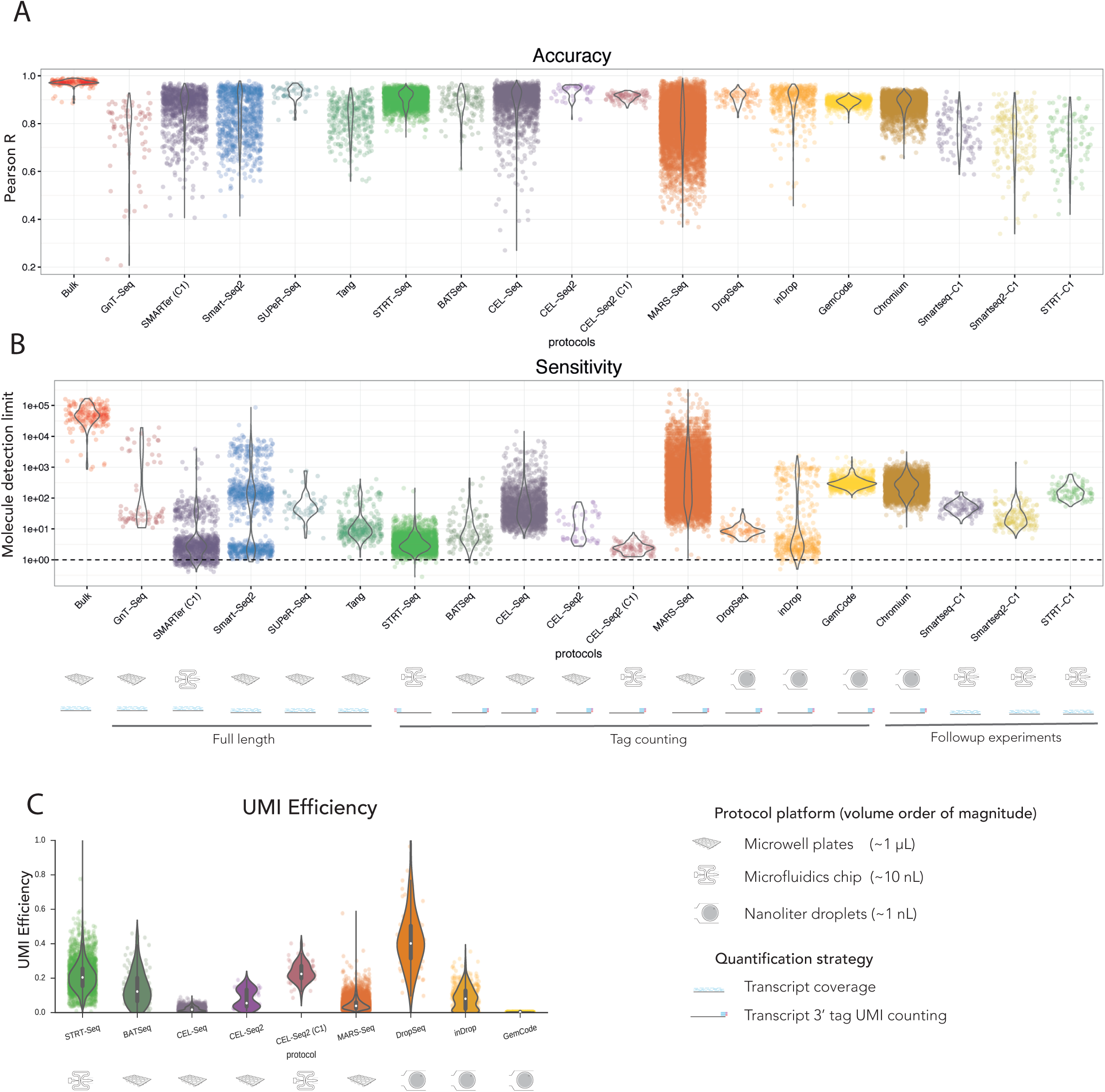
Comparison of performance metrics for different protocols. **(A) Accuracy.** While no protocol reaches the same level of accuracy as bulk RNA sequencing, in general accuracy is high regardless of protocol. Rarely is Pearson correlation lower than 0.6, though some protocols have longer tails with lower accuracy. This might be indicative of a heterogeneous success rate for the protocol. **(B) Sensitivity.** All protocols have greater sensitivity than bulk RNA-sequencing, indicating that the low-input problem they are trying to solve has indeed been addressed at some level. Sensitivity can vary greatly though, between protocols but also across samples within a protocol. Several protocols have the potential to measure single-digit spike-in molecules. **(C)** An alternative approach to measure sensitivity when assuming measured counts correspond to molecule counts is by the UMI efficiency. The efficiency is the number of counted molecules compared to the input number of molecules, *Ui* = *E* ∙ *Mi*. For the analysed UMI based data the UMI efficiency largely recapitulates the detection limit (Supp Figure 1B). However, note that the molecule counting assumption of UMIs does not hold perfectly (Supp Figure 2C-D).

It is also worth noting that the relationship between input molecules and measured expression (Figure 1D), can provide a guide as to the limit of accurate quantification on a per-cell basis. For instance, for the example cell shown in Figure 1D, quantification is extremely accurate down to about 10 molecules.

### Logistic regression to define technical spike-in sensitivity

We wanted to measure the sensitivity of each sample individually to be able to quantify variability in sensitivity for every method investigated, and to avoid biases due to uneven batch sizes. Simply using ratios of detected spike-ins at each abundance level would give poor resolution, because at most seven spike-ins share one abundance level. We elected to use a logistic regression model for detection in each sample, using detection of expression as the dependant variable. Our measure of sensitivity is the molecular spike-in input level where probability of detection reaches 50%.

When we apply logistic regression to all the public samples, we see in general that all the scRNA-seq protocols are more sensitive than regular bulk RNA-sequencing. In this respect, all protocols do resolve the low-input problem of single-cell experiments. We can also see that many protocols have the potential to sense as little as between one and ten input spike-in molecules. However, the amount of within-protocol variability for sensitivity is rather large, making it hard to rank the protocols based on this metric alone.

### UMI efficiency of tag-counting protocols

The majority of single cell RNA-sequencing protocols utilise a umi-tag counting strategy to achieve digital quantification of mRNA transcripts. Here one single tag from a single mRNA molecule is reverse transcribed, and gets an additional probabilistically unique random identifier sequence. These protocols create cDNA libraries with extremely low complexity, which could produce extreme amplification biases. The UMI on each tag should allow one to remove this, as it is added prior to amplification. The question then remains as to how efficient this entire procedure is.

The underlying assumption is that the number of UMIs of a gene U = E * M, where 0 < E < 1 (Supplemental Figure 2A). In this model E is the *UMI efficiency* and M is the number of transcript molecules of a gene. We fitted this model for every UMI-tag sample, and the results are presented in Figure 2C. These results recapitulate the results from the technical sensitivity analysis (excepting MARS-Seq data, Supplemental Figure 2B).

However, more in depth analysis shows this measure might not be as appropriate as it appears. If we extend the model to be U = E * M^c, the best fit should give values of the *molecular exponent* c close to 1 if the underlying UMI counting assumption is correct. Instead, we find that the best fit is systematically lower than 1 and with a mode of ~0.8 (Supplemental Figure 2C). This implies a saturation of UMI counts as a function of input molecules. This can be explained partially (but not fully) by differences in UMI length between the different protocols (Supplemental Figure 2D). For example, UMIs of length 4 can only count 256 unique molecules. In all it appears UMI counts do not perfectly reflect direct counts of molecules. Still, even in protocols with UMI’s as long as 10 base pairs, we observe the molecular exponent to be around 0.8 per sample on average.

### Endogenous transcripts are more efficiently captured than ERCC spike-ins

It is unclear to what extent the sensitivity and accuracy calculated based on exogenous spike-ins corresponds to the same metrics for endogenous mRNA. On the one hand, ERCC spike-ins have shorter poly-A tails than typical mRNA from mammalian cells^7^, making them harder to capture by poly-T priming. On the other hand, endogenous mRNA have intricate secondary structures and are bound to proteins, potentially reducing the efficiency of reverse transcription.

To investigate the relation between single-cell RNA-seq measurements of endogenous mRNA and ERCC spike-ins, we analysed single molecule fluorescent in-situ hybridization (smFISH) data and CEL-seq data from the same mESC line and culture conditions (Grun et al; smFISH molecule counts kindly provided by the authors). Based on smFISH data for 9 genes, CEL-Seq UMI counts correspond to 5-10% of smFISH counts. This is in contrast to the ERCC spike-ins in the same samples, where on average UMI counts correspond to 0.5-1% of input molecule counts (Supplemental Figure 2E).

Surprisingly, this data suggests that endogenous RNA is much more efficiently captured and amplified than ERCC spike-in molecules, at least in this data set. This implies that our sensitivity estimates above may be conservatives, and are likely to be underestimates rather than overestimates. (Note that the accuracy estimates would not be affected as they relate to relative concentrations of molecules, which will be preserved.) This difference in efficiency should be considered if for example one uses ERCC spike-ins to infer absolute molecule counts.

### Sensitivity is more dependent on sequencing depth than accuracy

The results of the per-sample accuracy and sensitivity analysis shows a large amount of within-protocol heterogeneity. Seeking to explain performance by technical factors, we find a relation with sequencing depth per sample. We used a linear model which considers a global effect of sequencing depth, including diminishing returns. The model infers an individual corrected performance parameter for each protocol. This way we can rank protocols from the public experiments accounting for the large technical factor of sequencing depth.

Accuracy is not strongly dependent on sequencing depth, supporting the theory above that heterogeneity of accuracy in a protocol relates to a general success rate of a protocol (Figure 3A). The best performing protocols in terms of accuracy are SUPeR-Seq, the only total-RNA single cell protocol to date, and CEL-Seq2, which uses In Vitro Transcription rather than PCR to amplify cDNA material.

**Figure 3.**
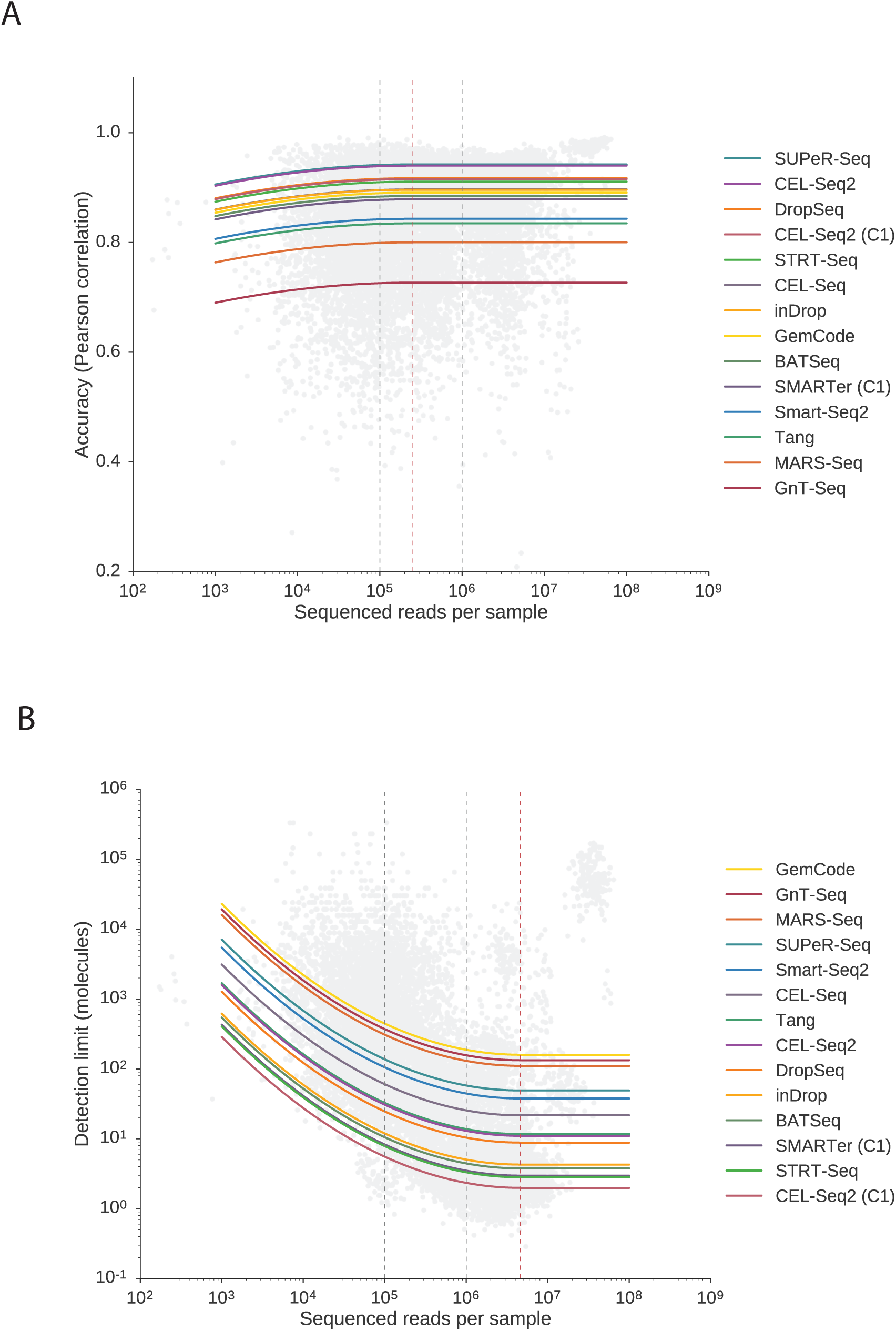
The effect of sequencing depth on performance metrics. We sought to investigate performance of the different protocols in light of sequencing depth, a major cost factor in single cell RNA-seq experiments. We modeled a global dependency on sequencing depth considering diminishing returns, with a distinct corrected performance parameter for each protocol. **(A) Accuracy** is only marginally dependent on sequencing depth. This means that even if you might not measure as many genes in a sample, the genes you do measure will be accurate. **(B) Sensitivity** on the other hand is critically dependent on sequencing depth. Comparing performance without accounting for sequencing depth is inappropriate, and th is explains the large amount of variability within protocol noticed before. The read level for saturation of sensitivity identified by the model is at 4.6 million reads per cell (dashed red line). We note that the improvement in sensitivity from 1 million reads is miniscule, and recommend this as a target depth for saturated single cell RNA-sequencing. In contrast, moving from 100 000 reads to 1 millions reads accounts for almost an order of magnitude better sensitivity. It is worth noting that not all assays require saturating detection, but care needs to be taken when comparing samples of different sequencing depths, even when using a compositional expression unit like TPM.

Technical sensitivity on the other hand is critically dependent on sequencing depth, and comparing sensitivity without accounting for differences in depth would be misleading (Figure 3B). With the corrected sensitivity score of the model, we can now rank the protocols. From the public data, the three protocols implemented in a C1 microfluidics system are the top three performing protocols in terms of sensitivity. The most recently developed of these, CEL-Seq2 (C1), was the most sensitive. We also note that the matched microwell plate implementation of CEL-Seq2 has worse sensitivity than the C1 implementation.

Since the model considers diminishing returns on the sequencing depth, we can identify when the performance metrics saturate from the model parameters. Accuracy already saturates at 250,000 reads, illustrating that it is not very dependent on sequencing depth in general.This means that as soon as one can detect expression, expression levels will be accurate and quantitatively meaningful.

Based on the model, sensitivity saturates at about 4.5 million reads per sample. We note though that the marginal increase from 1 million reads per sample to 4.5 million reads is miniscule, less than one fold change. At the same time, the marginal increase in sensitivity going from 100,000 reads per sample to 1 million reads per sample is almost an order of magnitude. Thus we recommend considering 1 million reads per sample as a good target for saturated gene detection.

It should be noted that not all studies need to saturate detection, since in many cases the genes of interest may be highly expressed. Another important note is that the sequencing depth is a technical feature, and we see here that the number of genes detected depend on this. When performing analysis of single cells, the technical sequencing depths of cells must be taken into account computationally, even for compositional expression units such as TPM.

### Degradation of spike-ins does not explain performance of experiments

The performance analysis of the scRNA-seq data inherently assumes the gold standard annotation of the spike-ins to be correct. However, RNA is fickle and can easily be degraded through the course of normal reagent handling. To quantify the effect of this, we performed an experiment where we used freeze-thaw cycles as a proxy for normal handling. Additionally, we allowed spike-ins to degrade to a level where they must be considered completely ruined, by letting them sit at either room temperature or at 37C overnight. During freeze-thaw cycles simulating normal handling, the effect on accuracy is miniscule, while for ruined spike-ins the accuracy difference is similar to that between protocols (Figure 4A).

**Figure 4.**
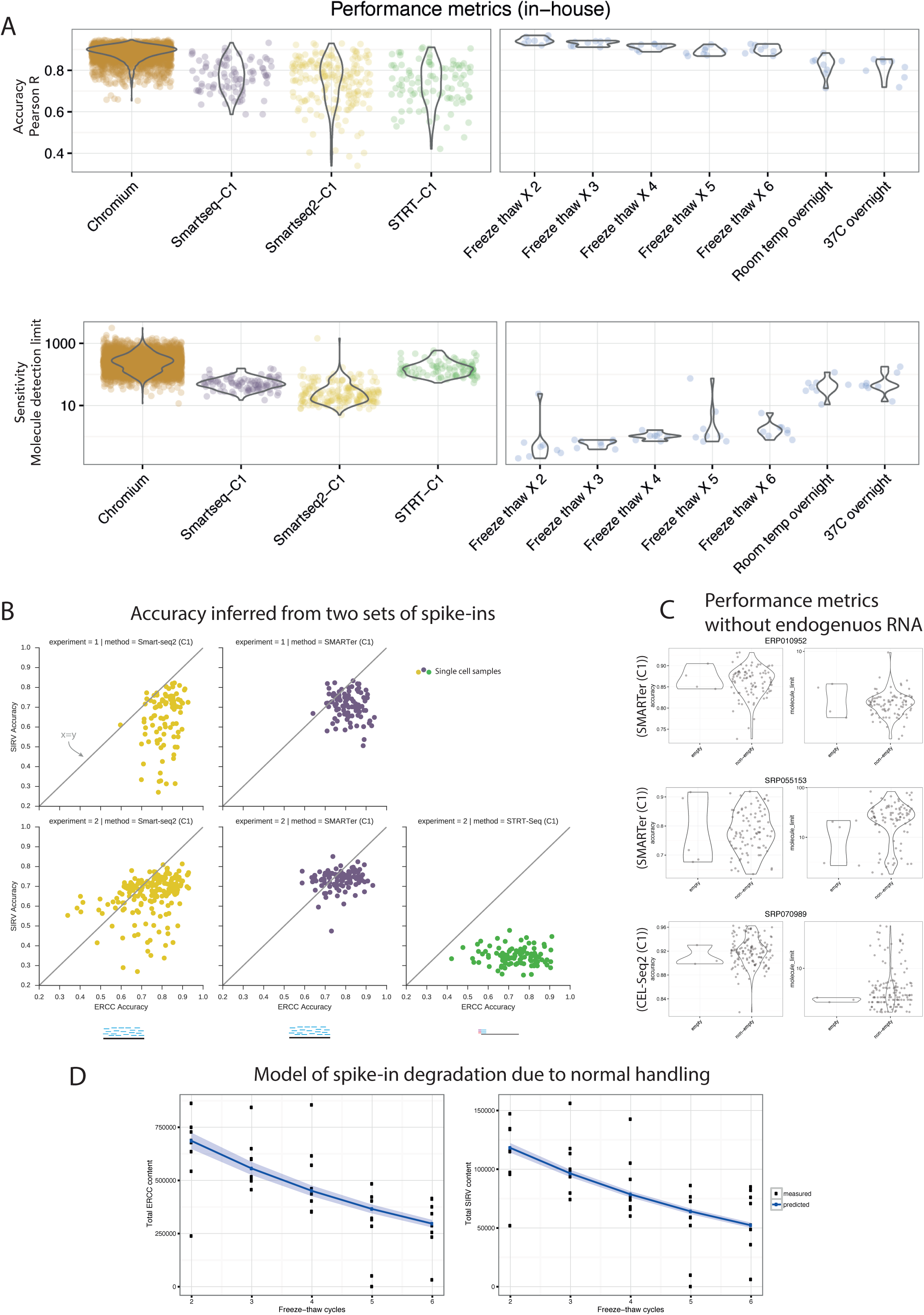
Investigation of factors affecting performance differences. **(A) Batch effects and RNA degradation**. While many of the protocols we investigated were replicated in different studies, it might be the case that performance of a protocol is confounded by laboratory competence, implementation platform, or similar external factors. To address this, we implemented three different protocols (SMARTer, Smart-seq2, and STRT-Seq) on the Fluidigm C1 platform, and performed these three in a single batch, using cells from the same culture as input. When performed on the same platform, we see an increased performance of Smart-seq2 over SMARTer. We wanted to assess how our performance metrics are affected by normal handling of reagents. For this, a series of freeze-thaw cycles were performed on spike-ins as a proxy for degradation through normal handling. Over the course of cycles, performance metrics did change, but to a lesser extent than the differences between protocols. When spike-ins are intentionally destroyed by being left overnight, the performance difference is on the same level as that between protocols. **(B) Different spike-in kits show same patterns of accuracy estimates.** In two experiments with matched conditions, we used two independent types of spike-ins (ERCCs and SIRVs) from two different manufacturers. Accuracy inferred from one set recapitulate accuracy from the other, indicating that manufacturer batch variability does not explain differences between protocols. Additionally, since SIRVs are different abundances of isoforms, this illustrates that, as expected, coverage based protocols can distinguish between isoforms while tag counting protocols cannot, as indicated by the result of the STRT-Seq condition. **(C) Endogenous RNA does not interfere with spike-in performance metrics**. The different studies we obtained the data from investigate different types of single cells, which have different amounts of endogenous RNA. To investigate whether a cell with a large amount of RNA would compete out the spike-in RNA and affect our performance metrics, we found three studies where empty wells (lacking endogenous RNA) were annotated. Comparing performance metrics of empty wells, where there is no competition with spike-ins, with non-empty wells, we see no improvement in any performance in empty well. This indicates that accuracy and spike-in sensitivity is not biased by endogenous RNA content in the samples. **(D) Relative spike-in abundance in a sample is affected by normal handling of spike-ins**. While we found our performance measures based on spike-ins are not affected by normal handling, comparing spike-in content in samples to endogenous RNA from mESCs, we noted significant changes in this ratio over freeze-thaw cycles of the spike-ins. This means inference of absolute amounts of mRNA in cells based on known input spike-ins needs to be aware of potential degradation and batch differences of spike-in amount. Fitting a Bayesian model of degradation, we found that a freeze-thaw cycle results in the loss of 19% of the spike-in RNA present in the sample. This was true both for ERCCs and SIRVs, suggesting this degradation rate is globally true for RNA in general, and should be taken into account when planning experiments with RNA.

When comparing samples in terms of technical sensitivity, degradation plays an even larger role. The assumption that the dilution provided is correct is what makes the metric comparable between experiments. Our results show that normal handling can only account for molecule limit difference within an order of magnitude, even for extreme cases such as six freeze-thaw cycles. Ruining the spike-ins by leaving them out overnight has a two orders of magnitude effect on the sensitivity metric (Figure 4A).

### During a freeze-thaw cycle, a sample loses a fifth of its RNA

It is common knowledge that freeze-thaw cycles degrade DNA and RNA. The absolute magnitude of this has so far not been quantified. In our investigation of spike-in degradation, we added healthy cells to the wells so that experiments would correspond to a normal single-cell RNA-seq experiment. This allowed us to compare the spike-in content with the endogenous RNA content in each sample, and compare to the number of freeze-thaw cycles.

Setting up a Bayesian model of RNA degradation with a degradation rate parameter p, we find, with very good confidence, that the degradation rate is 19% per freeze-thaw cycle. The degradation experiment also contained SIRV spike-ins (see next section below), and while measurements from these are more noisy due to mapping uncertainty, the order of RNA degradation is recapitulated when modelling these as well (degradation rate estimate of 18.5%). These values suggest a 20% degradation rate for freeze-thaw cycles as a robust estimate for RNA degradation globally during a normal experimental handling.

### SIRV spike-ins recapitulate accuracy results based on ERCC spike-ins

The Spike-in RNA Variant (SIRV) spike-in mix is a more recently designed set of spike-ins which are designed with RNA isoform studies in mind. The mix consists of 69 artificial transcripts which are produced to mimic the splicing patterns of 7 human genes. In Mix 2, these isoforms are input at four abundance levels. In two matched comparison experiments, we added both ERCC and SIRV spike-in mixes.

While four abundance levels are too few to give information about sensitivity, we can compare accuracy using both sets of spike-ins (Figure 4B). Generally, accuracy is systematically lower when using SIRVs, as expected because the ambiguous read mapping to the isoforms introduces a noise element. Overall, the pattern of relative accuracy based on SIRVs reflects that based on ERCCs in our SMARTer and Smart-Seq2 experiments. The SIRV spike-ins have a very poor accuracy in our STRT-Seq experiment, which is to be expected as the transcript tags alone cannot distinguish between different mRNA isoforms.

### Endogenous mRNA amount does not affect performance metrics based on spike-ins

When sequencing a cDNA library, the finite fragments generated are sampled from the pool of all cDNAs, so if the relative abundance of endogenous mRNA is higher, we are less likely to sample fragments from spike-ins. To verify that this was not a driving factor in our performance metrics, we investigated the published data to find experiments where empty and non-empty samples were both annotated in the same batch. We found three such data sets, and comparing accuracy and sensitivity between empty and non-empty samples shows that endogenous mRNA content does not affect our performance metrics (Figure 4C).

## Discussion

A previous study showed^8^ that read alignment to ERCC spike-ins varied widely between libraries and platforms, with some spike-ins having reproducibly poor behavior. This raises the question of how useful the spike-ins can be for calibration of absolute expression values. The ERCC spike-ins generally have very short poly-A tails of only 20 and to 26 bases long (the majority are 24 bases). In contrast, eukaryotic mRNAs have on the order of 250 base long poly-A tails^7^. Thus poly-T priming of ERCC spike-ins may be less efficient than for endogenous mRNA, leading all methods to appear somewhat less sensitive than they actually are for endogenous mRNAs. Furthermore, the ERCC spike-in RNA molecules are not capped at the 5’ end. This may lead to reduced efficiency of template switching compared with endogenous mRNAs^9^; template switching is required in several of the protocols. On the other hand, spike-in RNA does not have the RNA binding proteins or complicated secondary structure associated with endogenous RNA.

In order to compare these trade-offs between the spike-in and endogenous RNA, we used smFISH data as a gold standard for numbers of endogenous RNA molecules per cell, as compared to the calculated numbers of spike-in molecules per cell (Supplementary Figure 2E). This data suggests that endogenous RNA is more efficiently captured and amplified than spike in RNA by about one order of magnitude. This would lead all protocols to appear less sensitive than they actually are. Because of this, we must point out that while we use the “spike-in molecule detection limit” as a sensitivity measure, the ability to detect 10 spike-ins might not correspond to an ability to detect expression from 10 mRNAs. Nevertheless, the global ranking of the protocols remains relevant. These issues will not affect the estimates of accuracy, as all ERCC spike-ins in a sample will be equally affected.

Many of the protocols for single cell RNA-sequencing are extremely promising and provide tremendously powerful, high resolution techniques for unbiased genome-wide dissection of cell populations and their transcriptional regulation. Here, we show that while these methods vary widely in their detection sensitivity, with lower limits between 1 and 1.000 molecules per cell, their accuracy in quantification of gene expression are all very good.

While sensitivity depends critically on sequencing depth, accuracy is less linked to depth, and both features are linked closely to the molecular biological protocol used to generate the data. If lowly expressed genes are of interest in answering a particular biological question, a protocol with high sensitivity is more suitable, while for other applications, this may not matter. Protocols with low lower detection limit will also be able to provide insight into smaller differences of expression.

Overall, it appears that miniaturizing reaction volumes increases sensitivity, though implementations of this might make it harder to perform controlled experiments attempting to minimize batch effects. Regardless, sequencing more than around a million reads per sample provides a poor return on investment. In the future, improvements in protocols as well as decreases in the price of sequencing will further boost our ability to answer new questions in biology using single cell transcriptomics.

## Methods

### Mouse embryonic stem (mES) cells culture

Wildtype E14 mouse ES cells (kindly provided by Pentao Liu, Wellcome Trust Sanger Institute) were cultured on gelatin coated dishes using Knockout DMEM (#10829; Gibco), 15% Fetal Calf Serum (FB-1001/500; batch tested from Labtech), 1x Penicillin-Streptomycin-Glutamine (#10378-016; Gibco), 1x MEM NEAA (11140-035; Gibco), 2-mercaptoethanol (31350-010; Gibco) and 1000U Leukemia Inhibitory Factor (LIF; #ESG1107). Mycoplasma-free tested mES cells were passaged every 2-3 days.

### SMARTer, Smart-Seq2 and STRT-Seq on C1

E14 mESCs were trypsinized to obtain single cell suspension and passed through 30µm filter (CellTrics; #04-0042-2316) to remove debris and clumps. Cells were processed using the C1 Single Cell Auto Prep System (Fluidigm; #100-7000 and #100-6209) following the manufacturers protocol (#100-5950 B1). Briefly, we perform SMARTer, Smart-seq2 and STRT-Seq each across three small C1 Open App IFCs (5-10µm; #100- 5759). The specific sample preparation steps for the three protocols (SMARTer^10-14^, Smart-seq2^15^ and STRT-Seq^16-19^) were downloaded from Fluidigm Script hub. Dissociated single cells are loaded and captured on C1 Open App IFCs, followed by manually inspection serving as quality control measure to demarcate empty well, doublets or debris containing wells. We use two different spike-in RNA controls for batch-matched comparison of different protocols. 92 ERCC spike-ins (#4456740; Lot# 1411014; Ambion) and 69 SIRV spike-ins (#SKU025.03; E2 Spike-in RNA Variant Control Mixes; Lexogen) were mixed (0.5µl 1:500 diluted ERCCs + 0.6µl 1:500 diluted SIRVs) and added to respective Lysis buffer master mixes for SMARTer (20ml), Smart-Seq2 (27ml) and STRT-seq (20ml). 9ml of the respective Lysis master mix is added to each Open App C1 IFCs. The subsequent steps (Cell lysis, cDNA synthesis by reverse transcription and PCR reaction) are performed as described on Fluidigm Script hub.

### SMARTer and Smart-Seq2 on C1

Similar to above, E14 mESCs were trypsinized to obtain single cell suspension, passed through 30µm filter (CellTrics; #04-0042-2316) to remove debris and clumps. Single cell suspension was processed using SMARTer and Smart-seq2 in parallel across two C1 Single Cell Auto Prep System (Fluidigm; #100-7000 and #100-6209) following the manufacturers protocol (#100-5950 B1). Smart-seq2 protocol was downloaded from Fluidigm Script hub. The cells were loaded, captured on C1 Open App IFCs, followed by manually inspection. Both ERCC and SIRV spike-ins were mixed (0.5µl 1:500 diluted ERCCs + 0.6µl 1:500 diluted SIRVs) and added to respective Lysis buffer master mixes for SMARTer (20ml) and Smart-Seq2 (27ml). The subsequent steps (Cell lysis, cDNA synthesis by reverse transcription and PCR reaction) are performed as described on Fluidigm Script hub.

### Spike-in degradation experiment using Smart-Seq2 on plates

We used new tube of Spike-ins, ERCC (#4456740; Lot# 1412014; Ambion) and SIRV (E2 mix; #SKU025.03; Lot#216651530; Lexogen) for this experiment. Briefly, 1:100 dilutions of ERCCs and SIRVs are mixed together resulting in spike-in master mix (1:200 final dilution; termed ‘x2 Freeze-thaw’). The spike-in master mix is split into three tubes; one left overnight at 37°C (Condition 1), one left overnight at room temperature (Condition 2) and third kept overnight at −80°C. The following day the third tube (from −80°C) was subjected to multiple freeze-thaw cycle wherein the tube was thawed at room temperature for 2-5minutes, an aliquot was taken and re-freezed in dry ice. We repeated this freeze-thaw cycle an additional 5 times (Condition 3 to Condition 7). All the spike-in mixes (Condition 1-7) were subsequently diluting to a final 1:1000,000 dilution. A 96-well plate for Smart-seq2 was prepared by dispensing 2ml Smart-Seq2 lysis buffer (0.2%Triton, 1:20 RNAse inbhibitor, 10mM Oligo-dT_30_VN, 10mM dNTPs) across each well. 1µl of spike-in mix per condition (Condition 1-7) was added to each well column-wise such that each column represented a single condition with 8 replicate wells. E14 mESCs after removing debris and clumps using 30µm filter were FACS sorted (BD Influx; BD Biosciences) into 96-well plate. The first three wells (row-wise) across the 96-well plate were sorted to have matched bulk 500, 50 and 5 cells. The 96-well plate was immediately spun and frozen on dry-ice prior to Smart-seq2 protocol as described^15^.

### Library preparation and Sequencing

Representative cDNA from single cells across three C1 runs and Smart-Seq2 (on plates) was checked using High Sensitivity DNA chips using Bioanalyzer (5067-4626 and 5067-4627; Agilent Technologies). Single cell cDNA from SMARTer^10-14^ and Smart-Seq2 C1 IFCs and Smart-seq2 (on plates) was tagmented and pooled to make libraries using Illumina Nextera XT DNA sample preparation kit (Illumina; FC-131-1096) with 96 dual barcoded indices (Illumina; FC-131-1002). The clean up of library and sample pooling was performed using AMPure XP beads (Agencourt Biosciences; A63880). All protocols are described in the Fluidigm protocol (100-5950), Fluidigm Script Hub and Smart-seq2 protocol^15^. The STRT-Seq libraries were made and sequenced as previously described^16-19^. The Single cell libraries from SMARTer and Smart-Seq2 C1 IFCs and Smart-seq2 (on plates) was sequenced across 1 lane of HiSeq V4 (Illumina) using 75bp/125bp paired-end sequencing.

### 10x Genomics Chromium experiment

The Single Cell Gel Bead kit (#120217), Single cell chip kit (#120219) and Single cell library kit (#120218) were used along with 10x GemCode Single Cell Instrument as per manufacturer specifications and manuals (Document # CG00011; Revision B). An equal mix of Control Brain RNA (3µl; FirstChoice Human Brain Total RNA; #AM7962) and ERCC spikes (3µl 1:4 dilution; #4456653) was mixed to form ‘2x Control RNA+ERCC’ master mix. We further diluted ‘2x Control RNA+ERCC’ to ‘1x Control RNA+ERCC’ with PCR grade water. We made two single cell master mix preparation using 3µl of ‘2x Control RNA+ERCC’ and ‘1x Control RNA+ERCC’ respectively instead of single cell suspension (adjusted with 34.4 µl Nuclease-Free water). Rest of the steps were followed as per manufacturer's manual (Document # CG00011; Revision B). Each 10x library was sequenced across HiSeq2500 (2x lanes; Rapid Run) as per Wellcome Trust Sanger Institute sequencing guidelines.

### Data Sources

Raw read data from published studies was downloaded from either ENA or SRA, as listed with accession numbers in Table 1. Information regarding concentration and volume of ERCC mix in each sample was gathered from the original publications (also indicated in Table 1) or through direct communication with authors in ambiguous cases.

The expression table for mESC-STRT had non-standard names annotating the ERCC spike-ins, and through personal communication with the authors we were given a table for converting these to the names as provided by Life Technologies. Additionally we were informed by the authors that the final spike-in dilution noted as 1:50000 in Islam et al^18^ had in actuality been 1:20000.

The concentrations of the ERCC solution in the Dendritic-MARS table was ambiguous as there were two different values in the GEO table and in the text of the paper. Communication with the authors clarified that these referred to different volumes. The volume and dilution described in the GEO table was used. Thirty samples were excluded as they were annotated as not having had ERCC spike-ins added to them.

For the K562-SMART data it was unclear which of all data sets had actually used spike-ins, and personal communication with the authors provided the names of the two batches which had had spike-ins added.

A table with notes on individual data sets is provided (Table 1).

### RNA-Seq data processing

For coverage based data, relative abundances were quantified using Salmon 0.6.0, with library type parameter -l IU and the optional flag --biasCorrect. The Salmon transcriptome indices were built by adding ERCC sequences to cDNA sequences from Ensembl. For samples with mouse background, this was Ensembl 83 cDNA annotation of GRCm38.p4. For samples with human background, this was cDNA annotation from Ensembl 78 of GRCh38, and for samples with zebrafish background, the Ensembl 77 annotation of Zv9. Finally, for samples with frog background, this was Ensembl 84 annotation of JGI4.2.

In order to process all UMI-based data in a coherent way, we developed a quantification strategy based on pseudo-mapping, and counting up evidence for (transcript, UMI) pairs. We implemented this in a publicly available command line tool which we call ‘umis’. The tool as available at https://github.com/vals/umis/ as well as in the Python Package Index, and in Bioconda.

The principle is to transfer information from a (UMI, tag) pair to a (transcript, UMI) pair based on which transcript the tag maps to. Since UMI-based methods only use 3’ or 5’ end tags of cDNA, which can be as short as 25bp, mapping of these tags are commonly ambiguous. Our strategy for this is to weight a (UMI, tag) pair by the number of transcripts the tag maps to. After (UMI, tag) pairs were mapped with either RapMap or Kallisto in pseudobam mode, only (transcript, UMI) pairs with a user specified minimum amount of evidence are counted (default 1). This can be either on the gene or transcript level. In the 10x Genomics Chromium data we detected 70,000 and 45,000 droplets with respect to the samples. For the sake of computational memory efficiency we uniformly sampled 2000 droplets out of all detected droplets to count the umi tags per droplet.

### Analysis

An ERCC spike-in was considered detected when the estimated TPM of that ERCC was greater than zero. For UMI-based data, a spike-in is detected when at least one copy of an ERCC molecule is inferred.

The amount of input spike-in molecules for each spike, for each sample, in each experiment was calculated from the final concentration of ERCC spike-in mix in the sample.

Calculation of the accuracy of an individual sample was done by the Pearson correlation between input concentration of the spike-ins and the measured expression values. If less than 8 spike-ins were observed, the accuracy was set to infinity, as we consider this to be insufficient evidence to estimate the accuracy.

For the logistic regression model of each sample’s detection limit, the probability of detecting a spike-in at a given input level is modeled by the logistic function:

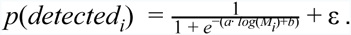

We used the LogisticRegression class from the linear_model module of the machine learning package scikit-learn^20^. The fit was performed with the liblinear solver and the optional argument fit_intercept=True. The logistic regression analysis was limited to samples with at least eight spike-ins detected. The detection limit was chosen as the molecular abundance where the logistic regression model passes 50% detection probability:

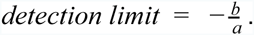

To investigate the UMI efficiency of UMI based protocols, we used a linear model where the only parameter was the efficiency:

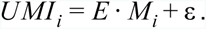

As we mention in the text though, the data fits a model much better where there is a non-one exponent parameter on the number of input molecules:

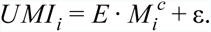

When we model the relation between read depth and performance metrics for individual protocols, we use a linear model with a quadratic term for read depth to capture diminishing returns on investment. The model considers the read depth effect to be global, and has a categorical performance parameter for each protocol:

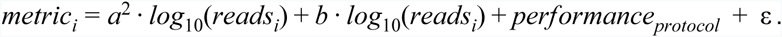

Here the performance metric will plateau and saturate when

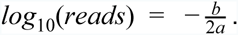

The linear models were fitted and analysed using the OLS regression function in the statsmodels Python package.

In the spike-in degradation model the degradation rate p and the cellular fraction F were inferred by a Bayesian approach using Stan (R package rstan v 2.10.1). The model was specified as the following: p was sampled from a uniform distribution between 0 and 1, F_i_ for each spike-in i was drawn from a normal distribution with mean 0.5 and standard deviation 1. F_ij_ was estimated by a normal distribution with mean F_i_ *(1-p)^j^, where j was the j-th freeze-thaw cycle and standard deviation sigma sampled from a uniform distribution between 0 and 20. The model was run with 5000 iteration steps, 1000 warm up steps and 4 chains.

All data needed for our analysis is provided as Supplemental Table 1.

## Acknowledgements

We are grateful to Oliver Stegle and Jong Kyoung Kim for helpful discussions and comments on the manuscript. We also wish to extend our gratitude to Sten Linnarsson and Amit Zeisel who gave invaluable support in implementing STRT-Seq in our lab. The study was supported by Cancer Research UK grant number C45041/A14953 to A.C. and C.L, European Research Council project 677501 - ZF_Blood to A.C. and a core support grant from the Wellcome Trust and MRC to the Wellcome Trust – Medical Research Council Cambridge Stem Cell Institute. ERC grant ThSWITCH to S.A.T. (grant no. 260507) and a Lister Institute Research Prize to S.A.T. K.N.N. was supported by the Wellcome Trust Strategic Award “Single cell genomics of mouse gastrulation”.

All RNA-seq expression data from our experiments are available with ArrayExpress accession XXXX.

## Supplemental Figure Legends

Table 1

**Summary of studies used for this comparison.** The column **Name** contains our identifier for the data set. In **Description of data** there is a short summary of the cells studied in each reference. The **Reference** column contains the original publication the data stems from. In **Protocol** the name of the protocol, and what section of transcripts they aim to capture is annotated. Protocols that use Unique Molecular Identifier barcodes to quantify expression are indicated in the **UMIs** column. The number of cells (or samples) are indicated in the **Cells / Samples** columns, and the number of batches these are divided into is noted in the **Batches** column. Finally, the **Accession ID** provides the ID used to retrieve the data from public databases. ID’s starting with SRP or ERP refer to the European Nucleotide Archive, while ID’s starting with GSE were retrieved from the Gene Expression Omnibus.

**Supplemental Figure 1.**
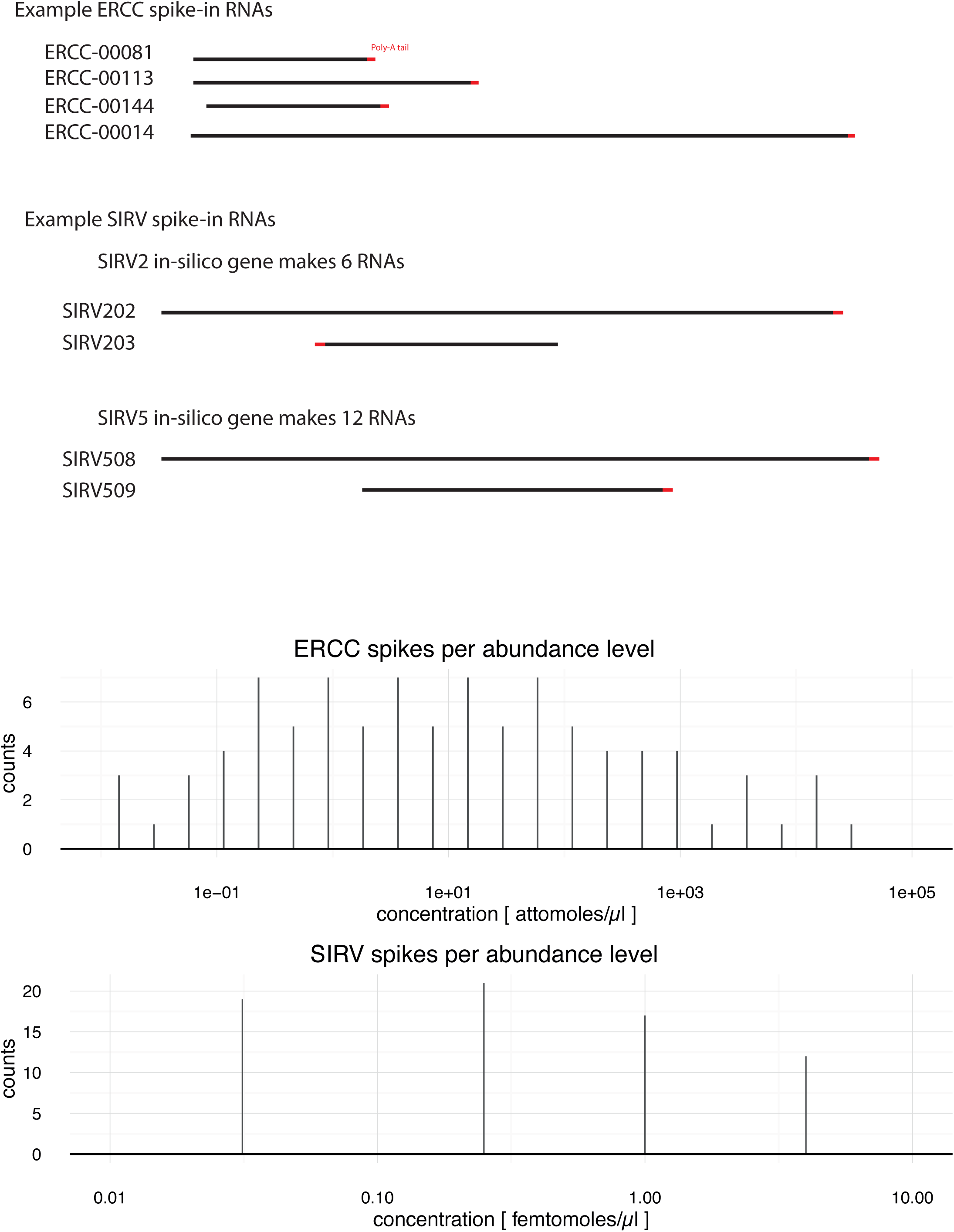
Comparison and overview of spike-in sets. ERCC spike-ins consist of 92 very distinct sequences based on bacterial genes logarithmically distributed across 22 abundance levels (in Mix 1), with poly-A tails ranging from 20 to 26 base pairs. SIRV spike-ins are 69 sequences, modeled after sequences and splicing patterns in 7 human genes. In Mix 2, which we used, the SIRV molecules are present at 4 abundance levels, with virtual alternative isoforms from each gene present at each abundance level. All SIRV molecules have 30 base pair long poly-A tails.

**Supplemental Figure 2.**
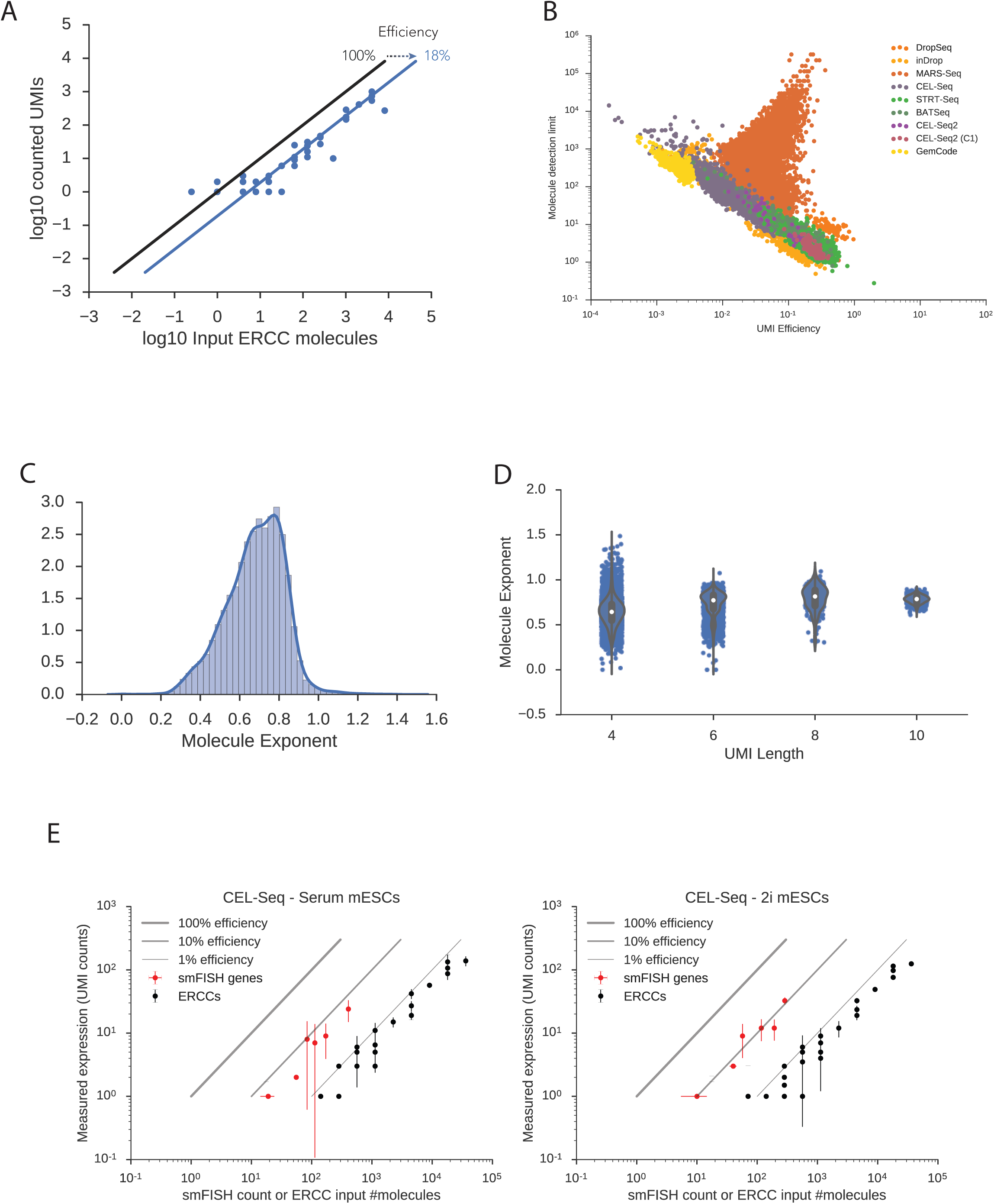
UMI efficiency as an alternative metric of sensitivity. **(A)** Assuming that UMI counts correspond to a count of the fraction of molecules successfully captured by the RNA-sequencing process, in log-log space the efficiency corresponds to the offset from perfect correspondence between input molecules and counted UMIs. **(B)** With the exception of data from the MARS-Seq protocol, spike-in detection limits correspond well with UMI efficiency measures. The spike-in detection limit can however also be used for coverage based data quantified by TPM. **(C)** The assumption with UMI counting as a quantitative measurement is that efficiency is the only factor determining differences between real counts and observed counts. However, fitting a model with a non-one exponent on the number of input molecules shows this is almost in all cases < 1. This means UMI counts underestimate expression of highly expressed genes. **(D)** The saturation of UMI counts can be partially explained by short UMIs. If an experiment uses too short UMIs, eventually the number of possible observable UMIs plateau. However, even for very long UMIs, such as 10 base pairs, the mean molecule exponent is 0.8, indicating some additional unexplained factor is causing a saturation of UMI counts. **(E)** Averaged efficiency comparison of endogenous genes and ERCC spike-ins. The data by Grun et al had smFISH measurements for 9 genes in the same experimental conditions as the single-cell RNA-seq data. Assuming 100% capture rate for smFISH, we can compare average smFISH counts with average UMI counts. Round markers correspond to median value across cells, and bars correspond to 95% confidence interval across cells. The smFISH counts suggest UMI counts for endogenous transcripts are on the order of 5-10% on average, while ERCC spike-in UMI counts correspond to 0.5-1% efficiency on average.

**Supplemental Figure 3.**
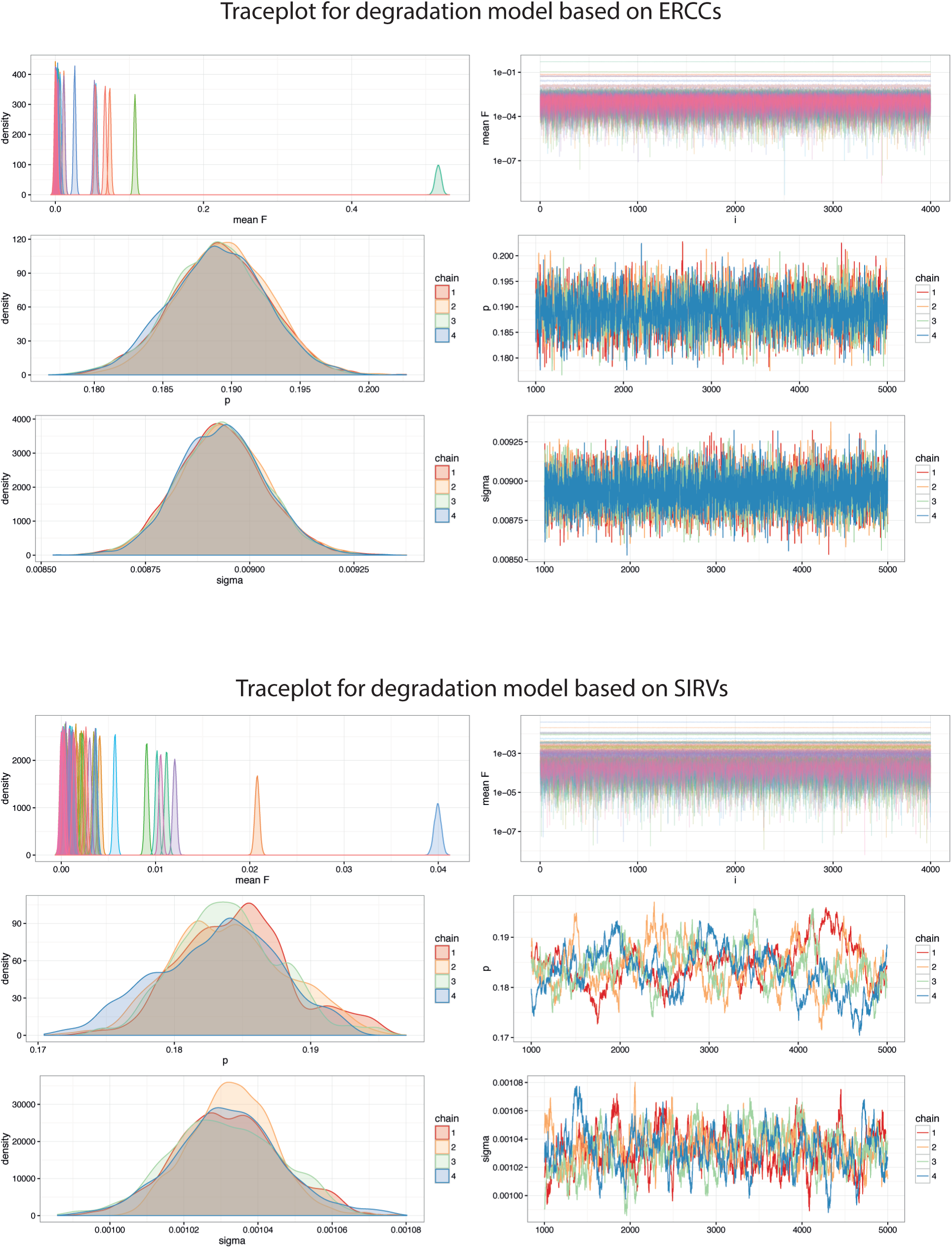
Traceplots from Bayesian models of degradation. The posterior samples from the model parameters in *stan^21^* for both the ERCC and SIRV analysis show very narrow confidence intervals and good correspondence between the different sampling chains. The SIRV based model is slightly noisier, which can be expected, as isoform-level expression when multiple isoforms are present is a harder quantification problem than quantifying expression of the unique ERCC sequences. For the ERCC model, the mode of the degradation rate parameter p is 19%, and for the SIRV model it is 18.5%.

## References

1. Macaulay, I. C. & Voet, T. Single cell genomics: advances and future perspectives. PLoS Genet. 10, e1004126 (2014).

2. Stegle, O., Teichmann, S. A. & Marioni, J. C. Computational and analytical challenges in single-cell transcriptomics. Nat. Rev. Genet. 16, 133–145 (2015).

3. External RNA Controls Consortium. Proposed methods for testing and selecting the ERCC external RNA controls. BMC Genomics 6, 150 (2005).

4. Jiang, L. et al. Synthetic spike-in standards for RNA-seq experiments. Genome Res. 21, 1543–1551 (2011).

5. Munro, S. A. et al. Assessing technical performance in differential gene expression experiments with external spike-in RNA control ratio mixtures. Nat. Commun. 5, 5125 (2014).

6. Hashimshony, T., Wagner, F., Sher, N. & Yanai, I. CEL-Seq: single-cell RNA-Seq by multiplexed linear amplification. Cell Rep. 2, 666–673 (2012).

7. Viphakone, N., Voisinet-Hakil, F. & Minvielle-Sebastia, L. Molecular dissection of mRNA poly(A) tail length control in yeast. Nucleic Acids Res. 36, 2418–2433 (2008).

8. SEQC/MAQC-III Consortium. A comprehensive assessment of RNA-seq accuracy, reproducibility and information content by the Sequencing Quality Control Consortium. Nat. Biotechnol. 32, 903–914 (2014).

9. Kapteyn, J., He, R., McDowell, E. T. & Gang, D. R. Incorporation of non-natural nucleotides into template-switching oligonucleotides reduces background and improves cDNA synthesis from very small RNA samples. BMC Genomics 11, 413 (2010).

10. Mahata, B. et al. Single-cell RNA sequencing reveals T helper cells synthesizing steroids de novo to contribute to immune homeostasis. Cell Rep. 7, 1130–1142 (2014).

11. Treutlein, B. et al. Reconstructing lineage hierarchies of the distal lung epithelium using single-cell RNA-seq. Nature 509, 371–375 (2014).

12. Wu, A. R. et al. Quantitative assessment of single-cell RNA-sequencing methods. Nat. Methods 11, 41–46 (2014).

13. Pollen, A. A. et al. Low-coverage single-cell mRNA sequencing reveals cellular heterogeneity and activated signaling pathways in developing cerebral cortex. Nat. Biotechnol. 32, 1053–1058 (2014).

14. Buettner, F. et al. Computational analysis of cell-to-cell heterogeneity in single-cell RNA-sequencing data reveals hidden subpopulations of cells. Nat. Biotechnol. (2015). doi:10.1038/nbt.3102

15. Picelli, S. et al. Smart-seq2 for sensitive full-length transcriptome profiling in single cells. Nat. Methods 10, 1096–1098 (2013).

16. Jaitin, D. A. et al. Massively parallel single-cell RNA-seq for marker-free decomposition of tissues into cell types. Science 343, 776–779 (2014).

17. Grün, D., Kester, L. & van Oudenaarden, A. Validation of noise models for single-cell transcriptomics. Nat. Methods 11, 637–640 (2014).

18. Islam, S. et al. Quantitative single-cell RNA-seq with unique molecular identifiers. Nat. Methods 11, 163–166 (2014).

19. Zeisel, A. et al. Brain structure. Cell types in the mouse cortex and hippocampus revealed by single-cell RNA-seq. Science 347, 1138–1142 (2015).

20. Pedregosa, F. and Varoquaux, G. and Gramfort, A. and Michel, V. and Thirion, B. and Grisel, O. and Blondel, M. and Prettenhofer, P. and Weiss, R. and Dubourg, V. and Vanderplas, J. and Passos, A. and Cournapeau, D. and Brucher, M. and Perrot, M. and Duchesnay, E. Scikit-learn: Machine Learning in Python. Journal of Machine Learning Research 12, 2825–2830 (2011).

21. Carpenter, B., Gelman, A., Hoffman, M., Lee, D. & Goodrich, B. Stan: A probabilistic programming language. J. Stat. Softw. (2016).

